# MapTurns: mapping the structure, H-bonding and contexts of beta turns in proteins

**DOI:** 10.1101/2024.01.02.573949

**Authors:** Nicholas E. Newell

## Abstract

**Motivation:** Beta turns are the most common type of secondary structure in proteins after alpha helices and beta sheets and play many key structural and functional roles. Turn backbone (BB) geometry has been classified at multiple levels of precision, but the current picture of side chain (SC) structure and interaction in turns is incomplete, because the distribution of SC conformations associated with each sequence motif has commonly been represented only by a static image of a single, typical structure for each turn BB geometry, and only motifs which specify single amino acids have been systematically investigated. Furthermore, no general evaluation has been made of the SC interactions between turns and the structures in their BB neighborhoods. Finally, the visualization and comparison of the wide range of turn conformations has been hampered by the almost exclusive characterization of turn structure in BB dihedral-angle (Ramachandran) space.

**Results:** This work introduces MapTurns, a web server for motif maps, which employ a turn-local Euclidean-space coordinate system and a global turn alignment to comprehensively map the distributions of BB and SC structure and H-bonding associated with sequence motifs in beta turns and their local BB contexts. Maps characterize many new SC motifs, provide detailed rationalizations of sequence preferences, and support mutational analysis and the general study of SC interactions, and they should prove useful in applications such as protein design.

**Availability and Implementation:** MapTurns is available at www.betaturn.com along with a broad, map-based survey of SC motifs in beta turns. HTML/Javascript code for a sample map is available at: https://github.com/nenewell/MapTurns/tree/main.

**Supplementary Information:** Supplementary File 1: Methods.

## 1 Introduction

Beta turns^1^, in which the protein backbone (BB) abruptly changes direction over four amino acid (AA) residues, represent the most common type of protein secondary structure after alpha helices and beta sheets and play key structural and functional roles; for an overview, see *Roles of beta turns in proteins* at www.betaturn.com. Previous work (see brief reviews in^2,3^) has developed beta-turn classification systems at multiple levels of precision in the Ramachandran space of a turn’s BB dihedral angles, partitioning the turns into classes defined by angular ranges (the “classical” turn types^4^) or derived by backbone clustering^2^, and sequence preferences in turns have been computed for these classes. Some of these preferences have been rationalized with reference to the intrinsic properties of the side chains (SCs) involved (e.g. hydrophilicity, common in turns because they often lie close to the solvent-accessible surface) or intra-turn interactions (commonly hydrogen bonds) in which the SCs take part, but the distribution of SC structures associated with each sequence motif has generally been represented at most by a single, “typical” conformation for each BB geometry, illustrated with a static image. Furthermore, there has been no systematic analysis of SC motifs in beta turns that involve more than one AA, or motifs that link turns to their local BB neighborhoods, likely stabilizing the turns and the structural motifs in which they are found. The current picture of SC structure in beta turns and their local contexts is therefore only a rough sketch.

## 2 Tool description

This work introduces MapTurns, a server for motif maps, which are 3D graphical, interactive conformational heatmaps of the BB and SC structure and H-bonding associated with individual sequence motifs in four-residue turns and their two-residue BB neighborhoods (their “tails", which represent turn contexts). Maps are produced not only for beta turns, but also for four-residue “strand” turns that are not included in the beta turn definition^2^ because one or both of their central residues lie in beta strands.

Maps for beta turns are generated from a redundancy-screened dataset of 102,192 turns by a three-stage, hierarchical clustering procedure which uses a Ramachandran-space BB clustering generated by a hybrid DBSCAN/k-medoids algorithm^2^ as its first stage. In the second clustering stage, the turns within each BB cluster which contain the motif are clustered by the conformations of the motif’s SC(s), using a k-medoids PAM algorithm^5^ in Euclidean space. Finally, in the third clustering stage, the N- and C-terminal tails of the turns within each SC cluster are independently clustered in Euclidean space using k-medoids PAM.

In the maps for strand turns, which are classified into three groups {E2, E3, E2E3} depending on which of the two central turn residues lie within a strand, BB clusters are derived from a new k-medoids PAM clustering in Ramachandran space (since strand turns were not included in the beta-turn BB clustering^2^), while SC and tail clusters are generated using the same methods applied for beta turns.

The map interface, which is presented across two web pages, reflects the clustering hierarchy: the upper-level screen (Figure 1) supports browsing the distributions of BB and SC structures associated with the motif, while the lower-level screen enables the exploration of the turn contexts (represented by the tail clusters) which are associated with the BB cluster selected on the upper-level screen, as well as the browsing and profiling of the underlying PDB^6^ structures in the dataset, aligned in Euclidean space^7^.

**Figure 1.**
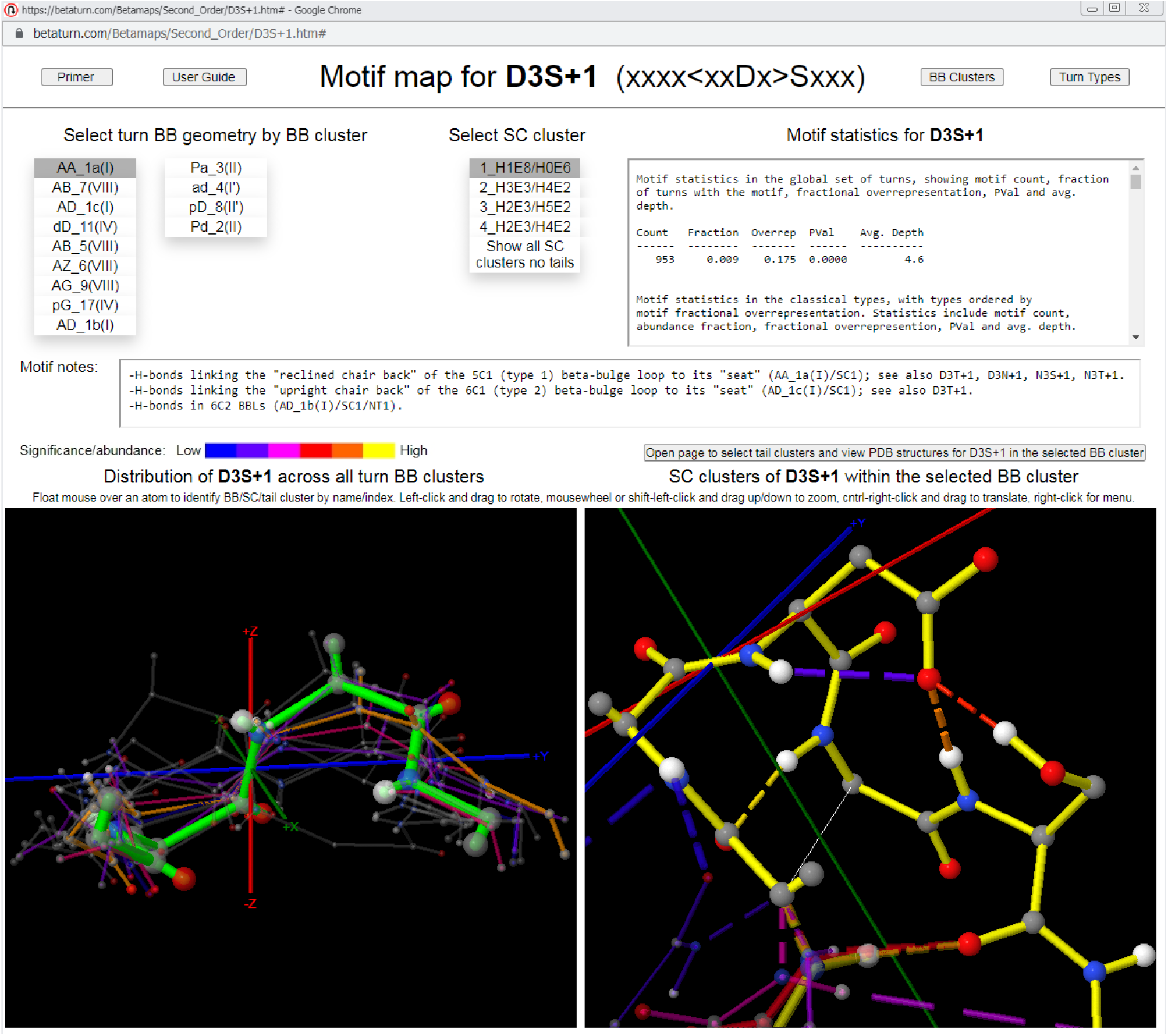
Upper-level motif map screen for the sequence motif that specifies Asp at turn position 3 and Ser just after the turn (D3S+1). The upper-level motif map screen supports the exploration of the recurrent BB and SC structures and H-bonds associated with a sequence motif in beta turns. BB and SC clusters are represented by their medoids, which are color-coded with a heatmap of relative motif statistical significance (for BB clusters, at left, with the selected cluster highlighted in green) or cluster size (for the SC clusters within the selected BB cluster, at right) and scaled according to cluster size. Navigation is achieved via menus that order clusters by statistical significance (for BB clusters) or cluster size (for SC clusters, in which all members contain the motif), and motif statistics are provided, along with text annotations for a selection of key motifs. In the figure, the D3S+1 motif’s most significant BB cluster is selected along with its largest SC cluster, and the SC cluster viewer at right shows that these choices are associated with SC/SC and SC/BB H-bonds linking the turn to the bulge in the common type 1 beta-bulge loop. The lower-level screen, opened by the button above the SC cluster viewer, supports the navigation of the contexts (tail clusters) associated with the motif within the BB cluster selected on the upper-level screen, as well as the browsing and profiling of the underlying PDB structures.

Clusters are displayed in JSmol^8^ viewers on each map page, represented by the structures of their medoids, which are color-coded as a heatmap of motif statistical significance (at the BB level) or cluster size (at the SC and tail levels, where all cluster members contain the motif), and scaled according to cluster size. At the SC level, the medoid structures convey the rotamer distribution of the motif’s SC(s).

Maps also display the frequency distributions of the SC/SC, SC/BB, and BB/BB hydrogen bonds associated with the motif, in the form of heat-mapped H-bonds connecting the corresponding atoms of the cluster medoids (see caveats in Supplementary Section 1.4).

Navigation in a map is achieved via menus listing the clusters at each level. Menu entries are ordered in the same way that medoids are heat-mapped - by the motif’s statistical significance in each cluster (for BB clusters) or cluster size (for SC and tail clusters), enabling the quick identification and browsing of important conformations at each level. Maps also incorporate embedded motif statistics, lists of the PDB addresses of the medoids and cluster members, and text annotations for a selection of notable motifs.

Motif maps are designed to support the rapid and intuitive exploration of the most important structures and interactions associated with a sequence motif. Extensive built-in help is provided, including text prompts, a primer with annotated map pages, and an in-depth user guide, as well as documentation for the beta-turn classification systems, which include the nine classical types^4^, eighteen BB clusters^2^, and 351 Ramachandran types^2,9,10^ in the dataset.

Motif-detection utilities available with MapTurns identify and rank the most statistically significant sequence motifs in each type, BB cluster or the global turn set. Motifs that are important in particular turn contexts, such as H-bonded structures (e.g. beta-hairpins^11^, beta-bulge loops^12,13^, alpha-beta loops^14^, helix caps^15,16^) or supersecondary structures in which a turn connects helices/strands, can be identified with the ExploreTurns^13^ tool, also available at www.betaturn.com.

A broad, map-based survey of structure and interaction in beta turns and their BB contexts is available with MapTurns, and serves as a “thumbnail index” to a selection of the most significant or otherwise notable SC motifs.

## 3 Implementation

Maps are implemented in HTML/CSS and Javascript, with embedded JSmol viewers. For best performance, browse with Chrome or Edge.

## 4 Applications

Motif maps provide a comprehensive picture of the distributions of structure and H-bonding associated with single-AA, pair, and many triplet sequence motifs in beta turns, and reveal the local BB contexts in which each motif is important and the roles the motif plays in those contexts. As the map-based survey shows, maps characterize many new SC motifs and support detailed structural rationalizations of sequence preferences, and the easy visualization and comparison of turn BB and SC conformations which maps provide should make them a useful tool for education as well as research.

Since beta turns adopt a broad range of BB geometries and exhibit a wide variety of SC interactions, including H-bonds, salt bridges and hydrophobic, aromatic-Pro, pi-stacking, pi-(peptide bond) and cation-pi interactions (see the motif survey), they can serve as a laboratory for the general study of these interactions, and maps support this work.

Maps also support mutational analysis, since a comparison of the maps for the motifs present in wild-type and mutant structures reveals the changes in structure and interaction that can occur with AA substitution(s). For example, a comparison of the map for the motif which specifies aspartic acid at the first turn position and arginine at the third position (D1R3, which forms one of the most significant salt bridges in beta turns), with the map for the motif which substitutes lysine at the third position (D1K3) shows the structural effect of the substitution. In BB cluster AA_1a(I), in which both motifs are most commonly found and short salt bridges are frequent within D1R3’s two largest SC clusters, the lysine SCs in D1K3’s largest SC clusters instead project away from D1, and the map shows no bridging in these clusters above the 20% display threshold.

The map also shows the structural compromises that must be made for salt bridging with D1K3: bridging can occur in the smallest BB cluster, but the Lys SC must bend sharply (the bridge H-bond appears unphysical in the map due to the ambiguity in H-atom labelling at the end of the Lys SC). The disruption of salt bridging in D1K3 compared to D1R3 is likely responsible for the motif’s much lower fractional overrepresentation in the BB cluster (10% compared to 59%).

Maps should prove generally useful in any application that can benefit from the comprehensive and detailed picture they provide of structure and interaction in turns and their contexts. One potential application is protein design, and in particular the design of binding and active sites, in which turns are common.

## Declarations

### Availability of data and materials

MapTurns is available at www.betaturn.com; for best performance, browse with Chrome or Edge. All data supporting the conclusions of this article are available from the Protein Data Bank at www.rcsb.org. HTML/Javascript code for a sample map is available at: https://github.com/nenewell/MapTurns/tree/main.

## Acknowledgements

The author wishes to thank Athena Newell for useful discussions and work as a research assistant and Kit Newell for work as a research assistant. The constructive suggestions and support of the attendees of ISMB/3DSig 2019 (Basel) and the 2019 and 2023 Annual Symposia of The Protein Society (Seattle and Boston) also contributed to the development of this project.

