## Supplementary information for "MapTurns: mapping the structure, H-bonding and contexts of beta turns in proteins"

### MapTurns

#### Supplementary information

### 1 | Methods

#### 1.1 Turn datasets

Beta turns are defined here as four-residue BB segments with a distance of no more than 7Å between the alpha carbons of their first and fourth residues and central residues with DSSP[1] codes outside the set {H, G, I, E} that specifies helix or strand[2]. Turn/tail structures were extracted from their PDB[3] files with the aid of a list of peptide chains with a maximum mutual identity of 25% obtained from PISCES[4]. Since sequence motif detection identified multiple motif "artifacts" generated by remaining local redundancy in the data, an additional step of turn-local redundancy screening was applied: turn/tail structures were clustered and filtered to reduce redundancy within five residues of turns to 40% or less; this threshold was chosen because it effectively reduced artifacts while preserving a dataset of sufficient size.

Turn structures were screened for quality by requiring that there be no missing residues within 5 residues of any turn and excluding structures with bond lengths that constituted outliers from established values. The final dataset of 102,192 turns was compiled with resolution and R-value cutoffs of 2.0Å and .25 respectively; average values in the dataset are 1.6Å and .20.

The three separate, smaller datasets of "strand" turns, for which one or more central residues are of DSSP type 'E', were generated with the same quality criteria as the beta-turn dataset. Maps for strand turns are labelled with the suffixes _E2, _E3, or _E2E3 indicating the residues that lie within strands; the corresponding turn datasets contain 5095, 3633 and 2982 structures respectively.

#### 1.2 Clustering

Motif maps are composed of medoids generated by three-stage hierarchical clustering. The first (BB) clustering stage uses a set of medoids generated from an ultra-high resolution beta-turn dataset using a hybrid DBSCAN/k-medoids procedure in Ramachandran space[2], modified slightly to improve the precision of map coverage by splitting the largest cluster, which contains turns with classical type I and type I-adjacent geometries, into three parts {AA_1a(I), AD_1b(I), AD_1c(I)}. Second-stage, SC clustering applies the k-medoids PAM algorithm[5] in Euclidean space to the SC and BB atoms of the amino acids specified by the motif in the turns which contain it in each BB cluster, identifying the most important recurrent SC conformations. Finally, third-stage (tail) clustering applies k-medoids PAM to the Euclidean-space coordinates of the BB atoms of the two-residue N- and C-tails of the turns in each SC cluster, identifying the most important recurrent contexts associated with a motif. Clusters are represented in the JSmol viewers by their medoids, or, when tails are displayed, by composites of medoids; see the primer and user guide for details.

Clustering in the datasets of E2-, E3- and E2E3-strand turns applies the same methods as those used for beta turns, except that the first-stage (BB) clusters are generated by a k-medoids PAM algorithm in Ramachandran space, with the number of clusters determined by a silhouette[6] analysis (this was necessary because strand turns were not included in the hybrid DBSCAN/k-medoids clustering procedure[2] used to classify beta-turn BBs).

Maps display structures in a turn-local coordinate system, established via a set of geometric definitions, which implicitly aligns the turns[7].

### 1.3 Computing sequence motif overrepresentation and p-value

Motif maps employ statistical tools to evaluate the over/under-representation and the significance of sequence motifs in types and clusters; the chi-square measure of significance is heat-mapped onto the BB medoids in the BB cluster view. The statistical methods[8,9,10] used to compute motif over/underrepresentation, chi-square and p-value are described in the map's user guide.

### 1.4 H-bond display

The distribution of SC and BB H-bond frequencies within and between clusters is displayed in a map in the form of heat-mapped dashed lines between the corresponding groups in the cluster medoids; frequencies from 20-100% are displayed using the color scale shown on the map. The criteria used to define H-bonds are given in the user guide. The heat-mapped "H-bonds" may appear unphysical due to differences in conformation between a cluster's medoid and its members or ambiguities in atomic labelling. Atomic labelling ambiguities may also produce inaccuracies in the displayed H-bond frequency distribution.

The set of H-bonds displayed by JSmol in PDB structures does not always exactly correspond to the H-bonds shown in the maps, due to differences in the H-bond definitions applied by MapTurns and JSmol (for example, JSmol does not apply separate cutoffs for the H-bond angles centered on the H atom and the acceptor atom).
